# Stronger bat predation and weaker environmental constraints predict longer moth tails

**DOI:** 10.1101/2024.05.05.590753

**Authors:** Juliette J. Rubin, Caitlin J. Campbell, Ana Paula S. Carvalho, Ryan A. St. Laurent, Gina I. Crespo, Taylor L. Pierson, Robert P. Guralnick, Akito Y. Kawahara

**Affiliations:** McGuire Center for Lepidoptera and Biodiversity, Florida Museum of Natural History, Gainesville, FL, USA; Department of Biology, University of Florida, Gainesville, FL, USA; Bat Conservation International, Austin, TX, USA; Department of Entomology, Smithsonian Institute, National Museum of Natural History, Washington, DC, USA; Florida Museum of Natural History, University of Florida, Gainesville, FL, USA

**Keywords:** Elaborate traits, predator-prey, Saturniidae, insectivorous bats, biogeography

## Abstract

Elaborate traits evolve via intense selective pressure, overpowering ecological constraints. Hindwing tails that thwart bat attack have repeatedly originated in moon moths (Saturniidae), with longer tails having greater anti-predator effect. Here, we take a macroevolutionary approach to evaluate the evolutionary balance between predation pressure and possible limiting environmental factors on tail elongation. To trace the evolution of tail length across time and space, we inferred a time-calibrated phylogeny of the entirely tailed moth group (*Actias + Argema*) and performed ancestral state reconstruction and biogeographical analyses. We generated metrics of predation via estimates of bat abundance from nearly 200 custom-built species distribution models and environmental metrics via estimates of bioclimatic variables associated with individual moth observations. To access community science data, we developed a novel method for measuring wing lengths from un-scaled photos. Integrating these data into phylogenetically-informed mixed models, we find a positive association between bat predation pressure and moth tail length and body size, and a negative association between environmental factors and these morphological traits. Regions with more insectivorous bats and more consistent temperatures tend to host longer-tailed moths. Our study provides insight into tradeoffs between biotic selective pressures and abiotic constraints that shape elaborate traits across the tree-of-life.

## 1. Introduction

Elaborate traits (complex, conspicuous derivations of pre-existing traits that serve a novel function [1]) provide a lens through which we can investigate opposing evolutionary pressures, as they are most likely to have emerged via strong selection. From the Narwal’s tusk [2] to the peacock’s train [3] to the porcupine’s quills [4], elaborate traits often play a role in high-stakes inter or intraspecific interactions – either to win potential mates or to evade potential predators. Due to their complexity and conspicuousness, these traits are commonly assumed to come with tradeoffs [5]. In some cases, tradeoffs have been empirically shown [6] but in many systems they can be hard to measure [7,8]. Frequently, when attempting to uncover tradeoffs, tests focus on short-term “acute tradeoffs” (i.e., increased energy expenditure, reduced maneuverability, etc. [5]). It can also be difficult to estimate these acute costs, given that traits evolve as integrated components of an animal’s biology and thus commonly occur in tandem with cost-reducing characteristics [9]. As a result, longer-term tradeoffs are usually the more relevant constraining force on trait elaboration [5]. Here, we use macroevolutionary analyses to investigate the relative roles of biotic and abiotic factors on the evolution of an elaborate wing trait in moths.

Moths in the family Saturniidae typically live for only a week as adults, during which time they do not feed and must locate mates at night to reproduce [10] while avoiding echolocating bats. At least five saturniid lineages have independently evolved hindwing tails with twisted and cupped ends [11]. Live bat-moth battles have revealed that these tails function as an anti-bat strategy. Experimental alteration, as well as natural variation of tail length in the luna moth (*Actias luna*) and the African moon moth (*Argema mimosae*), showed that as tail length increases, moth escape success also increases [11,12]. Compared with individuals whose tails were removed, those with tails got away >25% more from bat attack, despite there being no measurable difference in moth flight kinematics between treatments [12]. Tails therefore represent a powerful countermeasure to a nearly ubiquitous nocturnal selective force [13] and their success is highlighted by their repeated convergence across the saturniid family tree [12].

Studies on alternative pressures of hindwing tails have thus far been unable to uncover another driver or acute tradeoff. Mating trials using the luna moth have found no evidence that tails are used in mate selection [1]. Experimental studies with tailed and non-tailed luna moth models and diurnally foraging birds indicated that tails do not increase roosting moth conspicuousness to these predators, nor do they protect the moth by breaking search image [14]. These wing appendages also do not seem to be either a hindrance or an asset to evasive flight maneuvers based on in-battle kinematic analysis [12].

Tails may instead be limited by longer-term tradeoffs. In general, Lepidoptera wings grow proportionally with body size and both attributes are influenced by nutrition [15,16]. The longer amount of time a lepidopteran can stay in its larval form acquiring resources, the larger its body and traits are likely to be. Developmental studies testing tradeoffs between appendages in larval and pupal butterflies also indicate that growing and shaping wings has resource allocation costs [17,18]. An evo-devo study with the sphingid moth (the sister family to saturniids [19,20]) *Manduca sexta* showed that an increase in body size comes with a compensatory increase in development time or growth rate for wings to achieve appropriate allometric scaling [21]. Thus, seasonality is expected to lead to a broad pattern where adult lepidopteran body size and associated traits are smaller in more seasonally variable environments (generally higher latitude regions) and larger in more consistent (lower latitude) environments with longer growing seasons [22–26]. Insects therefore do not seem to conform to the same ecogeographic laws that has been ascribed to endotherms. That is, body size does not necessarily increase at higher latitudes (Bergmann’s Law) [24,27,28], nor do appendages (wings) appear to shorten at higher latitudes (Allen’s Law) [29]. Instead, body size and wing lengths are likely governed by other physiological forces. In the case of elaborate wing structures, it may be that the energetics of building extra wing material for a tail is a limiting factor for moths living in more seasonally variable environments with shorter growing seasons.

To test the macroevolutionary pressures that have shaped the elaborate hindwing tail trait, we focused our analyses to an entirely tailed clade of Saturniidae: *Actias + Argema*. This group is primarily distributed across Asia, from present-day Russia to Indonesia, and Africa [30], thus covering a broad range of habitat with many environmental conditions and exhibiting an array of hindwing tail lengths. We hypothesized that across their distribution and evolutionary history, large insectivorous bats have exerted a positive selective force on saturniid hindwing tails, but that elongation has been constrained by abiotic environmental factors. We further hypothesized that the association between bat predation pressure and moth body size has not been as strong as the association between bats and hindwing tail length, but that body size has been similarly susceptible to environmental constraints.

To test these hypotheses, we first built a well-sampled, time-calibrated phylogeny of the tailed moon moth clade (*Actias* + allies) and used this tree to trace the evolution of hindwing tails. In order to access the greatest possible number of observations, we employed a novel trait measurement approach where we extracted wing lengths from both digital museum collections images and community science photos on iNaturalist, using the moth’s antenna as a substitute scale bar, and verified this method using scaled museum images. We propose this approach for future researchers to use as a solution for the lack of scale bar in lepidopteran community science photos. To parse hindwing length trends from changes in overall size, we used moth forewing length as a proxy for body size [31,32]. We then investigated the effect of bat predation pressure (inferred abundance of sufficiently large insectivorous bat species) and environmental factors (mean annual temperature, average seasonal temperature variation, length of growing season, mean annual precipitation, and latitude) on our wing lengths of interest using phylogenetically-informed regressions. We predicted that in general, moth species whose distributions overlapped areas with greater insectivorous bat predation pressure would have longer tails than species inhabiting less bat-rich areas. We also predicted that this trend would be curtailed in regions with high seasonal temperature variability and thus more limited host plant growing season lengths. While we expected body size to follow similar patterns, we predicted the relationship between bat predation and size would be less pronounced. Our biologically-informed macroevolutionary approach provides a useful framework for scientists to examine the environmental and biological pressures driving trait elaboration across diverse taxa.

## 2. Materials and methods

### (a) Taxon sampling and DNA extraction

To reconstruct a well-sampled phylogeny of *Actias*, we used a combination of previously published data (7 ingroup species) [12] and newly sampled specimens from the McGuire Center of Lepidoptera and Biodiversity at the Florida Museum of Natural History (MGCL), Gainesville FL, USA (14 ingroup species; see Dataset S1 on Dryad for more details). We note that three of our newly sequenced specimens were from species that had previously been sequenced and published in [12] (*A. gnoma, A. selene, A. sinensis*). For the purposes of this study, we chose to sequence three new specimens, as the previous specimens did not have detailed enough locality data to include in our dataset. Our outgroup species were selected for their use as secondary calibration points in our phylogeny and came from sequences published in Kawahara et al. [20], as this analysis is the most comprehensive, fossil-calibrated phylogeny of Lepidoptera to date. We extracted DNA from both frozen, papered specimens (i.e., stored in an envelope in a -80 freezer since collection) and dried, pinned specimens (i.e., traditional museum preservation method) using an OmniPrep Genomic DNA Extraction Kit (G-Biosciences, St. Louis, MO) and evaluated DNA quality using agarose gel electrophoresis and quantity using Qubit 2.0 fluorometer (ThermoFisher Scientific). We sent our extracts to RAPiD Genomics (Gainesville, FL, USA) for library preparation, hybridization enrichment and sequencing using an Illumina HiSeq 2500 (PE100).

We analyzed our dataset using the Anchored Hybrid Enrichment (AHE) pipeline of Breinholt et al. [33]. We direct readers to this paper for detailed methods, but in brief, this pipeline uses an iterative probe-baited assembly process to clean raw reads and return an aligned set of orthologs for each locus in the probe kit. Because saturniids are in the superfamily Bombycoidea, we used the Bom1 probe kit (895 total loci) with *Bombyx mori* as our reference taxon [19]. We focused our analyses to coding regions (exons) and used MAFFT to align our sequences. We removed all loci that had <50% taxon coverage, leading to a total data set of 535 nuclear loci (40% of possible loci). To ensure that each locus was in the correct frame and did not contain any spurious nucleotides, we visualized each file in AliView [34] and made any necessary manual edits. To assemble our supermatrices, we used FASconCAT-G v1.02 [35]. Cleaned probe regions and supermatrices can be found on Dryad.

### (b) Phylogeny and estimation of divergence times

We reconstructed phylogenies with maximum likelihood (ML) and Bayesian inference (BI) optimality criteria, in IQ-TREE v. 2.0.3 [36] and BEAST v. 1.10.4 [37], respectively (Figs. S1-3). We inferred our maximum likelihood tree using IQ-TREE with the following commands: for one tree we used the ‘MFP+MERGE’ model, which maximizes model fit by sequentially merging pairs of genes (Fig. S1). For best compatibility with BEAST, we also inferred an ML tree using ‘-m TESTMERGE’, which operates similarly to ParitionFinder [38], and then specified only BEAST-applicable models: JC69, TN93, K80, F81, HKY, SYM, TIM, TVM, TVMef, GTR. For both ML trees, we calculated support values using 1000 ultrafast bootstrap replicates via ‘-bb 1000’ and 1000 Shimodaira-Hasegawa approximate likelihood ratio test replicates via ‘-alrt 1000’. We used the ‘-bnni’ command, which reduces the likelihood of overestimating branch supports by employing a hill-climbing nearest neighbor interchange (NNI) technique [36] (Fig. S2). We also performed a multispecies coalescent analysis with ASTRAL-III (v. 5.7.5) [39], which infers a summary species tree from the individual loci files generated in the IQ-TREE analysis (Fig. S4). We used all the default settings for the Astral analysis and assessed branch support using local posterior probabilities where anything <0.95 is considered weak support. This tree did not conflict significantly with our ML tree and we focus our analyses to the ML and Bayesian trees.

To infer our BEAST trees, we used BEAUTI v.1.8.4 [37] to create our command file. To infer divergence times, we used four secondary calibration points from Kawahara et al. [20]: Lasiocampoidea/Bombycoidea + other leps (78.61 – 99.27 mya), Lasiocampoidea + Bombycoidea (74.15 – 94.4 mya), Sphinigdae + Saturniidae (56.86 – 75.42 mya), Saturniidae (33.82 – 51.24 mya), Saturniini (14.54 – 30.63 mya). We constrained these calibration nodes with uniform distributions to stay within the age ranges inferred by [20]. To take advantage of the flexibility that BEAST offers regarding branch evolution rate, we used an uncorrelated relaxed clock model and drew from the lognormal distribution at each branch [40]. For our nucleotide substitution rate models, we used the substitution model, base frequencies, and site heterogeneity models identified by ModelFinder in IQ-TREE for each partition (23 partitions). We used our phylogeny inferred by IQ-TREE as the starting tree (Fig. S1), with calibration nodes manually set within the bounds of their age ranges using Mesquite [41]. We built Bayesian trees with either a fixed tree topology, to constrain the tree to the topology of the maximum likelihood input, or a classic operator mix, to allow BEAST to infer topology. To compare different models of evolution, we used “path sampling/stepping-stone sampling” marginal likelihood estimates (MLE) to determine whether a Birth-Death (constant rate of speciation and extinction applied) or Yule (special case of Birth-Death where extinction is null) prior best fit our data (Table S1) [42]. We performed separate runs that varied by operator mix and tree priors for 200 million generations each, sampling every 20000. All analyses were performed on the University of Florida’s high-performance computing cluster, HiperGator2.

### (c) Ancestral range estimation

To estimate ancestral ranges, we used the R package BioGeoBEARS [43] in RStudio (v. 2022.12.0+353) to fit a dispersal-extinction-cladogenesis (DEC) model. Under the DEC model, region occupancy is allowed to change along branches for each species via range expansion or reduction. Region occupancy can change at nodes via region-specific speciation where either both daughter species inhabit a range, one daughter species inhabits a subset of the range and the other inhabits the larger multi-area range, or the daughters split the ancestral range [44]. We used our BEAST maximum clade credibility tree and pruned outgroups to focus solely on species of interest and a few most closely related sister taxa. Our information about extant distributions of tailed moon moth species came from GBIF, iNaturalist (Research Grade only), and museum collection locality data, as well as expert input (Stefan Naumann and R.A.S.; see Supplementary Archive 4 on Dryad). Following Toussaint & Balke [45] and Lohman et al. [46], we defined seven regions based on biogeographical patterns and barriers (e.g., oceans, mountains): Africa (F), Americas (A), Europe (E), Philippines (H), Indomalaya + Greater Sunda Islands (M), East Palearctic (P), and Wallacea (W) (Figs. 1, S5). We built a dispersal multipliers matrix following Toussaint et al. [47]. According to this schema, probabilities of dispersal are penalized by the number of land masses that the animal must travel through to make it to another land mass or the size of the dividing body of water. For example, the dispersal probability from Wallacea to the Europe (Western Palearctic) is lower than from Wallacea to the Philippines (Tables S2-3, and see Supplementary Archive 3 on Dryad). This is a relatively young clade, and therefore has almost exclusively existed in a world of modern biogeographical configuration, however we did institute two time stratification layers to account for the closing of the Bering land bridge ∼5 mya [48,49]. While there were subsequent re-emergences of a Beringa bridge, the crossing likely would have been too cold for saturniid moths to use during the glacial maximum [49,50]. Thus, our time strata were set as 20 – 5 mya and 5 – 0 mya, with the only difference between them being a higher dispersal multiplier from East Palearctic to North America in the older time stratum (Table S3). We conducted two separate analyses, one more permissive and one more restrictive. Our permissive analysis allowed a maximum of 4 possible range outcomes, with nonadjacent ranges disallowed. To limit the number of permutations, and given that extant species exist in a maximum of two of our defined regions, our second analysis restricted possible range outcomes to 2 and defined the combination of regions that were possible (i.e., only adjacent regions).

**Figure 1.**
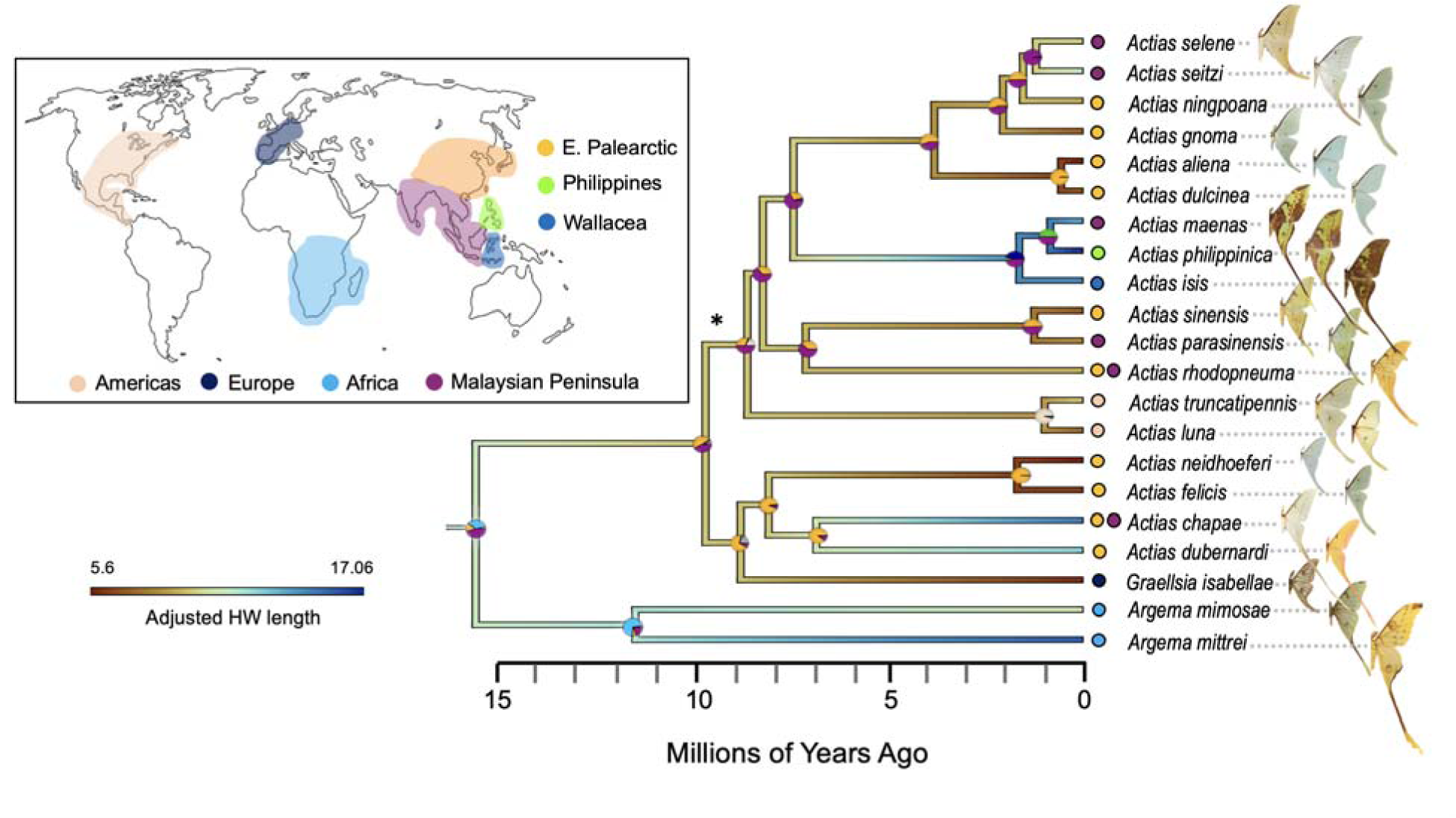
A time-calibrated tree of tailed moon moths (*Actias + Argema*) showing the inferred evolutionary and biogeographic history of long tails. Branches are colored by adjusted hindwing length (HW length/(Antenna length/mean species antenna length)) from images with a scale bar, with bluer colors representing longer hindwing tails and redder colors representing shorter tails. Median ages (in millions of years) were derived from a BEAST tree built with a Birth-Death prior using nodal calibrations from Kawahara et al. [20]. All support values from the starting maximum likelihood tree were 100/100, except at the node indicated by the asterisk, which was 80/100 (UFBoot/SH-aLRT). Colored circles represent probabilities of inferred ancestral ranges from our biogeographical (BioGeoBEARS) analysis, with colors reflecting the colored regions of the map at left.

### (d) Bat predation pressure

We generated species distribution models (SDMs) for bats carefully selected to represent likely saturniid predators. Our selection process identified bats that are primarily insectivorous and of sufficient size to be common predators that would exert strong evolutionary pressure on the moths, avoiding those that might occasionally pursue insect prey under limited circumstances (e.g., frugivores that may opportunistically prey upon insects, such as Phyllostomids [51]). We first identified all bat families where ≥ 50% of genera are aerial insectivores (18 out of 20 families) [52]. From these families we selected genera where ≥ 50% of species are sufficiently large (≥10g on average) aerial insectivores whose ranges overlap our moth species of interest (30 out of 129 genera), and finally filtered the data set to just species that also followed this description (Dataset S2). We chose this size threshold based on observations of bat behavior in the lab [11,12] and the general scaling of bat size to size of prey items [53]. After filtering, 179 species (59% of our initial target list) had sufficient occurrence records to reliably fit SDMs.

We leveraged an SDM-generation pipeline optimized for generating the distribution models of hundreds of species [54,55], customized to enhance performance for bats. First, we retrieved all bat occurrence records from the Global Biodiversity Information Facility (https://www.gbif.org/) and iDigBio (https://www.idigbio.org). We harmonized the taxonomy of records to the target species list using species definitions and synonyms from [52]. This matching was done using a manually-generated synonyms list for each targeted species. Data were cleaned with the “CoordinateCleaner” R package [56] and vetted with two rounds of manual checks. For each species with occurrence records, we defined accessible areas using a dynamic alpha hull encompassing cleaned points and buffered by 200 km. Dynamic spatial thinning was conducted on points as in [55], with the total accessible area determining the degree and rigor of thinning; model outputs were further tuned with manual checks to remove additional spatial biases.

We selected model predictors based on other macroecological studies of bats [57–60]; for example, we used topological ruggedness and roughness as proxies for cave and carst roosting habitats used by bat species [57]. Initial models were fit using 15 candidate predictors from BioClim (BIO1–2,4–6,12–17 [61]), three topographic (elevation, roughness, and terrain ruggedness index [62]), and one from MODIS data (percent tree cover [63]). The initial candidate predictors were selected to reduce collinearity while representing biologically plausible factors related to bat ecology. We further reduced model collinearity by iteratively refitting MAXENT models using default settings until all variance inflation factors were below 5. Using species-specific selected predictors, we quantitatively evaluated a suite of Maxent models with different tuning parameters to minimize model complexity and prevent overfitting using the R package ENMeval [64], using possible tuning parameters described in [55]. Final models (Table S4) were subject to an additional round of manual checks and adjusted or discarded if excessive commission or omission were apparent. We estimated species richness as the summed clog-log probability values from continuous surfaces, as recommended in [65]. To generate our estimates of bat abundance, we multiplied the clog-log probability SDM surface for each species by the population estimates provided in [66] and then divided by the sum of the clog-log surface to estimate the number of individual bats of a species in each 4664 m x 4664 m grid cell. Population estimates were generated via a regression model that incorporates average body mass (log transformed and z-score normalized) and IUCN Red List category for the species [66]. We then summed each species-specific density surface to estimate total bat density at each relevant location, and finally report the bat density surface in areas that overlap the moth species of interest (Fig. 2). For both bat richness and abundance, we extracted values at each site associated with moth hindwing length measurements. Code used to generate bat SDMs can be found on the Zenodo repository.

**Figure 2.**
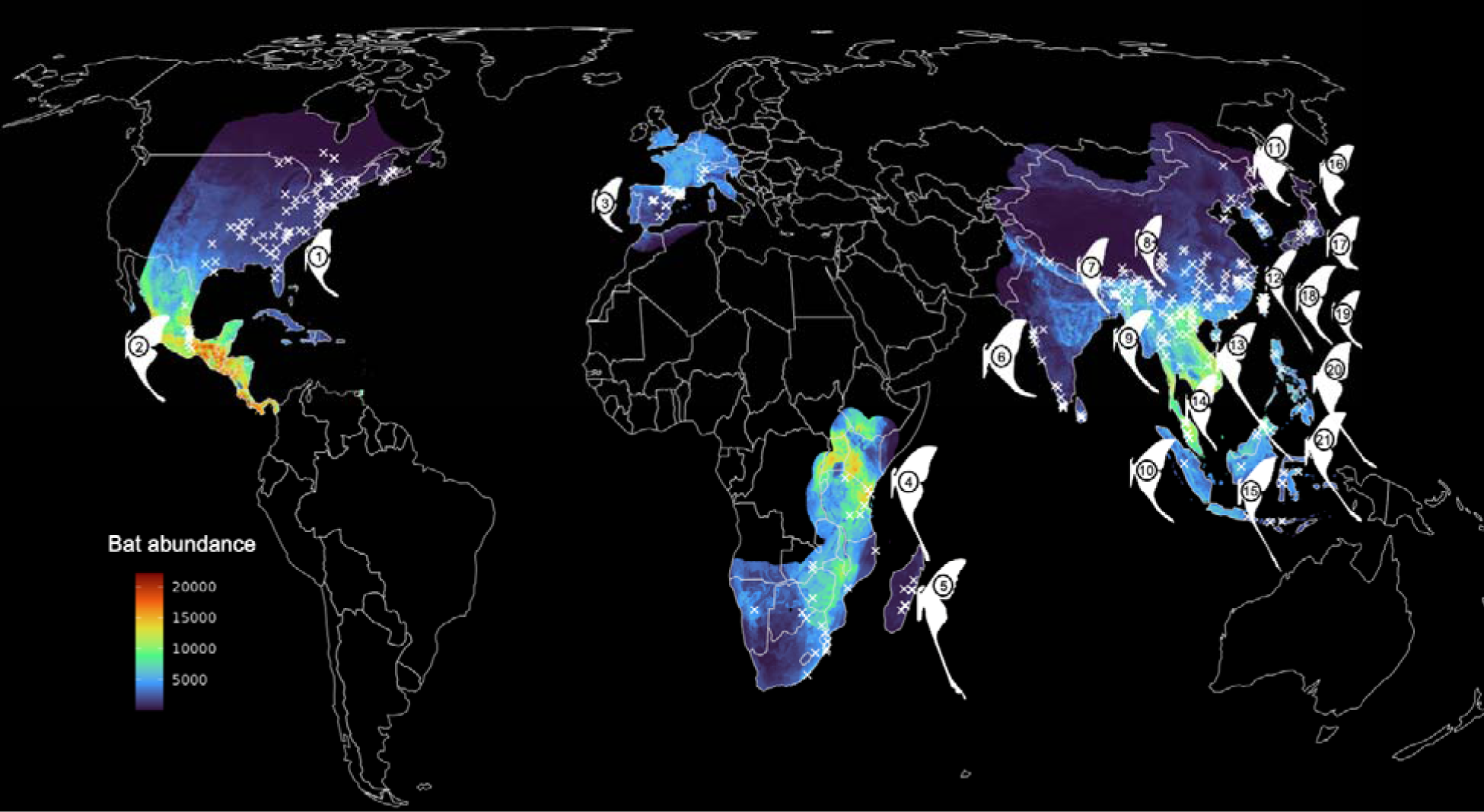
World map, pseudo-colored by bat predation pressure (sufficiently large, insectivorous bat abundance). We limited the visualization of our spatially explicit bat abundance estimates to only those areas that overlap with moth species of interest. Moth half-silhouettes indicate the general region where the species occurs. White x marks indicate precise moth observation locations, taken from museum collections, GBIF, and iNaturalist. We extracted environmental variable values from WorldClim and length of growing season values from the UN FAO FGGD LGP map for each moth point. We used these parameters, as well as measurements from the associated moth photo and estimates of bat abundance at each point, to build our phylogenetically-informed models. Species names: 1) *Actias luna,* 2) *A. truncatipennis,* 3) *A.* [*Graellsia*] *isabellae,* 4) *Argema mimosae,* 5) *Argema mittrei,* 6) *Actias selene,* 7) *A. parasinensis,* 8) *A. felicis*, 9) *A. rhodopneuma*, 10) *A. seitzi*, 11) *A. dulcinea*, 12) *A. dubernardi,* 13) *A. chapae,* 14) *A. sinensis,* 15) *A. maenas,* 16) *A. gnoma,* 17) *A. aliena,* 18) *A. ningpoana,* 19) *A. neidhoeferi,* 20) *A. philippinica,* 21) *A. isis*.

### (e) Hindwing tail trait acquisition

To extract wing measurements and associate these measurements with the individual’s coordinates, we gathered photographs from both natural history collections, including MGCL, American Museum of Natural History (AMNH), New York, NY, USA, and Stefan Naumann’s collection, (Berlin, DE), and publicly sourced data repositories, including GBIF and iNaturalist (see Dryad for all photos). To scrape images and associated coordinates from these online repositories, we used the function *occ_cite* in the R package “rgbif” [67] (see script on Dryad). While iNaturalist vastly increased our number of observations per species over museum specimens alone, photos on this site are unstandardized and most often are not associated with a scale bar. We therefore sought to find an alternative means of extracting a measurement of tail length. We determined that the most commonly visible components of the moth in these pictures were the forewings, hindwings, top-half of the thorax and antennae. In bat-moth interactions, the distance between the moth’s body and tail tips are most predictive of escape success [12]. We therefore were most interested in absolute tail length for our analyses. As a result, our goal was to find a component of the moth’s body that was as agnostic to body size as possible, to be used as a relatively standardized scale bar. We found that antennae length, unlike thorax width, had a low correlation with FW length (body size) in these species (Fig. S6). To further standardize this metric, we found the mean antenna length for each species and divided the individual antenna measurements by their species average. We then divided the forewing and hindwing lengths of each individual by their adjusted antenna length. To verify that these adjusted wing lengths led to similar absolute wing measurements in both scaled (museum) and unscaled (iNaturalist) photos, we performed Wilcoxon ranked sum tests for the forewings and hindwings of all species (see Fig. S7 legend for results). We visualized the overlap (mean ± SD) between adjusted wing length measurements from scaled photos and non-scaled photos and verified that both of these overlapped the “true” absolute wing measurements from scaled photos (Fig. S7). We further verified this approach by comparing true wing lengths from a subset of calibrated photos (with a scale bar) with their adjusted wing lengths (omitting the scale bar). That is, for this latter analysis, the comparison is between the exact same photos to test this method. Wilcoxon ranked sum tests again revealed no significant difference between true wing lengths (with a scale bar) and adjusted wing lengths (without a scale bar) (Table S5). While this approach leads to a slight underestimation of wing lengths for moths whose antenna are larger than their species mean, and a slight overestimation of wing lengths for moths whose antenna are shorter than their species mean, these differences are not statistically significant (Table S5) and the relationship between forewing and hindwing length within an individual remains quite consistent, as these wing lengths are relatively tightly correlated (r^2^ = 0.60). To ensure that our results for different species were not biased by the number of scaled or unscaled photos that were available for it, we used the adjusted wing length measurements for all of our analyses. All measurements were extracted using imageJ v.1.53t [68], via the “segmented line” tool to get antenna length from top to junction with the head and the “straight line” tool to get wing and thorax width lengths. Hindwing length was from tip of the tail to junction with thorax and forewing length was from tip of the forewing to junction with thorax. Thorax width was measured as the width of the prothorax (Fig. S6). When possible, we took measurements from the right side of the moth’s body, however, we would use the left side when elements from the right were unavailable or were less planar than from the left. We used only male moths for all analyses, as they are better represented in both collections and community scientist repositories and likely face higher bat predation due to flying more to locate females [69]. We also made sure all measurements taken from publicly sourced images were of high-quality, and of relatively un-damaged specimens and that the camera was at a perpendicular angle to the animal, to prevent inaccurate measurements due to distortion.

### (f) Phylogenetically-informed trait analysis and ancestral state reconstruction

To determine the strength of biotic and abiotic pressures on relative hindwing length, we used the function *pglmm* from the R package “phyr”, a mixed model approach to estimate evolutionary phenomena, accounting for phylogeny and spatial correlation [70]. We used adjusted hindwing length (HW length/(Antenna length/mean species antenna length))) as our response variable. To create our abiotic predictor variables, we extracted bioclimatic and growing period data for each moth occurrence in our dataset. Climatic variables came from the historical WorldClim dataset (2.5 arc-minute resolution, via the R package “raster”), which averages values between the years 1970-2000 (https://www.worldclim.org/data/bioclim.html). We extracted length of growing period (LGP) values from The Food and Agriculture Organization Food Insecurity, Poverty and Environment Global GIS Database (UN FAO FGGD) [71] using ArcGIS Pro v.2.6.6 and found the median of each LGP range. The LGP is determined by soil temperature and available moisture, accounting for transpiration, for crop growth. From these various data sources, we had a total of six predictors of interest in our models: bat abundance (described above), mean annual temperature (°C; code from Worldclim: BIO1), seasonality (standard deviation of mean annual temperature*100; BIO4), average annual precipitation (mm; BIO12), and median length of growing period (days). The phylogenetic covariation matrix and moth species were set as random effects in our models to account for relationships between the species and multiple occurrences per species. All variables were mean center scaled using the R function *scale* to make them comparable across highly varying units of measurement. To ensure that variables were not multicollinear (vif < 3 [72]), we used the *vif* function from the R package “car”. We ran a series of models with single or multiple predictors and used their DIC scores from the pglmm regression to determine best fit (Table S5 and in Supplementary Archive 1 on Dryad). While we felt it was important to include latitude in our models, due to its relevance in many other macroevolutionary studies, its posterior distribution was very broad, overlapped the zero line in all models, and it uniformly increased the DIC scores of our models. It did not change the relationships between our response variables and other parameters, however. We therefore maintain it in our models but do not discuss it as an important predictor (Fig. 3; and see code and outputs in Dryad).

**Figure 3.**
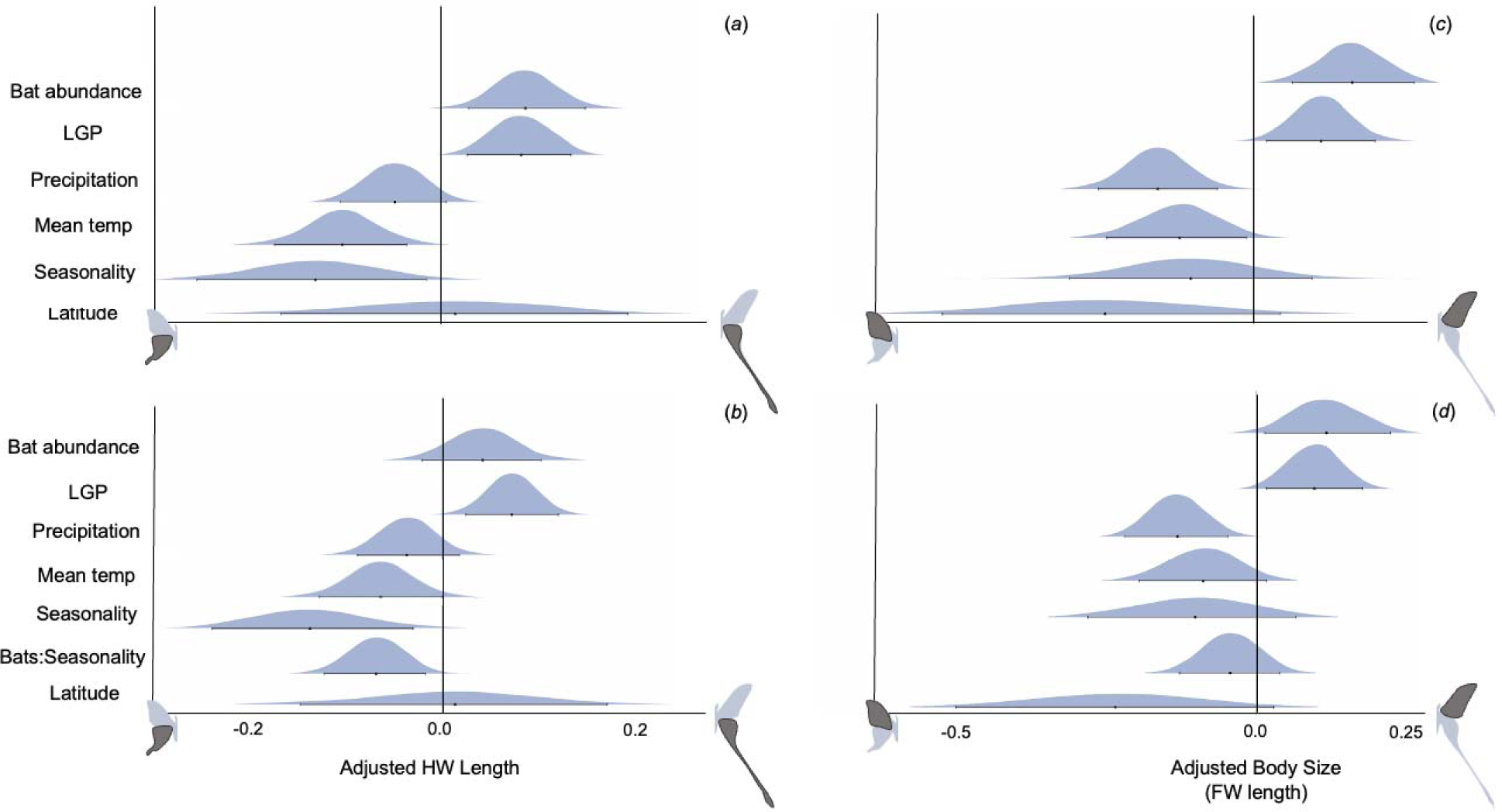
Tail length and body size in the tailed moon moth group are positively related to bat predation pressure and constrained by environmental factors. We find that insectivorous bat abundance is positively correlated with (a) hindwing tail length and (c) body size (forewing length) in *Actias* + *Argema* (Saturniidae). Length of growing period (LGP) is also positively correlated. Average annual precipitation, average annual temperature, and seasonality (standard deviation of temperature across the year) are negatively correlated with both tail length and body size. Thus, areas with higher bat abundance and longer periods of plant productivity are associated with longer tailed moth species. When we include an interaction term between bat abundance and seasonal temperature variation (bat:seasonality) for (b) tails and (d) body size, we find that some of the power is removed from bats as a driver of tail length. Although this parameter overlaps the zero line, there is still an ∼0.90 probability that bats have a positive relationship with tail length. This indicates that while bat abundance and seasonality have their own relationship with each other, they both still have independent effects on moth tails. Central tendency dots indicate parameter estimates and error bars are 95% credible intervals from the best fit phylogenetically-informed linear regression analyses. All predictor variables are mean center-scaled to make them comparable across units. Adjusted hindwing and forewing lengths are wing length/(antenna length/mean species antenna length).

We conducted comparative trait analyses and ancestral state reconstruction using the *ContMap* function and estimated phylogenetic signal using the *phylosig* function in the R “Phytools” package [73]. While we used only scaled museum specimens for this analysis, we generated the same metric that we used for ecological models to maintain consistency: HW length/(Antenna length/mean species antenna length)). To determine whether the best-fitting evolutionary parameter underlying this trait evolution was Brownian motion (BM; random walk), Ornstein-Uhlenbeck (OU; adaptive peaks), or early burst (EB; rapid then slow morphological evolution), we used the R “Rphylopars” package [74]. We used SURFACE [75] in R to test for convergent trait regimes across the phylogeny (see code and outputs on Dryad).

## 3. Results

### a) Phylogeny and estimation of divergence times

We built a 21-ingroup species AHE tree, including 14 newly sequenced specimens and seven previously sequenced specimens (see Dataset S2 for source and preservation type for each species). This represents about half of the total species in this group (40 species [76]), however, we accomplished broad sampling across the genus and the majority of missing species are in species complexes with those that we have represented in this tree. As with other phylogenetic studies of Saturniidae (e.g., 10, 11), we recovered a well-supported monophyletic group comprising *Argema* (*Actias + Graellsia*), sister to the Australian/Papua New Guinea clade *Syntherata* (*Opodiphthera* + *Neodiphthera*). Based on our log marginal likelihood comparisons (Table S1), we decided to use the Bayesian fixed tree with a Birth-Death model for all further analyses and interpretation (Fig. S3). We found that the *Actias* + allies diverged from these sister taxa ∼22 mya (HPD: 17.01 – 28.26 mya) (Fig. 1). While *Graellsia* has been known to be nested within *Actias* [77], and this was confirmed in our study (divergence from the other *Actias* in its clade ∼9 mya, HPD: 5.99 – 10.59 mya), we maintain the convention of using the *Graellsia isabellae* nomenclature. Within the *Actias +* allies ingroup, our tree largely agreed with the typology of these previous, less densely sampled AHE trees, as well as a study that reconstructed a phylogeny based on 16 *Actias* + allies species based on molecules, morphology, and behavior [78].

We found that Brownian Motion was the best fitting evolutionary model underlying the hindwing trait. That is, the evolution of tail length can be best described by a random walk, in comparison to a model with adaptive peaks or an early burst model (see scripts on Dryad). In line with this result, we found significant phylogenetic signal in hindwing length (where greater phylogenetic signal is represented by a K value closer to 1; Blomberg’s K=0.78, p= 0.002), and our SURFACE analysis [75] revealed only one hindwing length regime shift at the stem of *Argema + Actias* from the non-tailed sister taxa (Fig. S8). While we did not detect a signal of adaptive peaks in our dataset, our ancestral state reconstruction (ASR) analyses indicate that tails have repeatedly elongated in at least three separate lineages: *Argema mittrei + A. mimosae, A. chapae,* and *Actias maenas + A. philippinica + A. isis.* We also find evidence of tail length shortening in an equal number of lineages: *Graellsia isabellae, Actias neidhoeferi* + *A. felicis,* and *A. aliena + A. dulcinea* (Fig. 1). For comparative trait analyses using absolute hindwing length and absolute forewing length, see Figs. S9-10.

### b) Ancestral range estimation

To examine whether biogeographical history could explain some of the variation in hindwing tail length, we used BioGeoBEARS [43] to estimate ancestral ranges. Our 4-area range analysis resulted in unlikely combinations of ranges (Fig. S5A) and thus interpret the 2-area range analysis moving forward (Figs. 1 and S5B). We inferred a 0.83 probability that Indomalaya is part of the ancestral range of the tailed moon moths (including *Argema* and *Actias* species) and a 0.62 probability that Africa is in the ancestral range. Based on the inferred ancestral range of the common ancestor between *Actias* + allies and their closest sister taxa (0.76 probability Australia), we think it likely that the ancestral *Actias* moved from Australia to Malaysia by ∼15mya (HPD: 12.03 – 18.77 mya), after which point the *Actias* were extinct East of Lydekker’s line. The lineage leading to *Argema* split off from the rest of *Actias* and made it to Africa by about 11mya (0.97 probability)*. Actias* then diverged into a Palearctic group and an Indomalaya group by ∼9.5mya (HPD: 6.07 – 11.45 mya). It appears that there was a second wave of *Actias* movement into the Palearctic region by about 5 mya, leading to the extant short-tail species *A. dulcinea, aliena,* and *gnoma.* The diversification of *Actias* species across Wallacea and the Philippines islands ∼4 mya (HPD: 2.40 – 5.04 mya) likely originated from populations in Malaysia (0.91 probability). It is unclear how *Actias* arrived in the Philippines, as there is a roughly equal likelihood that they colonized this region via Wallacea or from the Indomalaya region. We also do not have strong inference as to the exact manner in which *Actias* colonized Northern America and Europe, but our analysis indicates that they did so from the Eastern Palearctic region, with lineages leading to *A. luna* and *truncatipennis* likely using the Bering Land Bridge (Figs. 1, S5).

### c) Phylogenetically-informed linear mixed models

Our global phylogenetically-controlled linear regression model (pglmm) revealed that hindwing length exhibits a positive relationship with bat predation pressure (parameter estimate (PE) of bat abundance: 0.082, 95% credible interval (CI): 0.028 – 0.137) (Fig. 3A). We also found a positive association between median growing season period and hindwing tail length (PE: 0.082, CI: 0.034 – 0.132). The credible interval for mean annual precipitation overlapped zero, but the probability that this parameter had a negative relationship with tail length was 0.94 and thus we interpret this along with the other environmental variables (PE: -0.045, CI: -0.097 – 0.007). Average annual temperature and annual seasonal temperature variation also displayed negative relationships with hindwing length, with seasonal temperature variation having the greatest effect size (PE seasonal temp variation: -0.126, CI: -0.236 to -0.070, PE avg temp: - 0.100 CI: -0.166 to -0.039) (Fig. 3A). We found that this global model performed better than a null model (which only accounts for phylogenetic relationships) and a model that contained all abiotic variables and excluded bat abundance, and performed slightly worse than an interaction model between bat abundance and seasonal temperature variation (DIC full model: 350, DIC null: 373, DIC no bats: 357, DIC interaction: 346). When we include the interaction, we find the same relationships between our parameters and hindwing tail length as in our global model. Under this framework, the bat parameter crosses the zero line, however there is still a 0.91 probability that bat abundance has a positive relationship with hindwing tail length (Fig. 3B). We found that while phylogenetic relationships alone explain much of the variance in hindwing length (r^2^ = 0.87), the global model explained more (r^2^ = 0.89). Additionally, removing bat abundance from the model decreased the explanatory power of the model by ∼2% and including the interaction term increased explanatory power by ∼1%, compared to the global model (*r2_pred* [79]; see code and outputs on Dryad). Breaking the dataset down by moth species also demonstrated that hindwing length was positively correlated with bat abundance in almost all species and was negatively correlated with seasonal temperature variation in almost all species (Fig. S11; see Table S6 for model structures).

The global pglmm analysis on forewing length (a proxy for body size [31,80]) showed similar relationships with all parameters (PE bat abundance: 0.149, CI: 0.065 – 0.233, PE LGP: 0.106, CI: 0.028 – 0.183, PE precipitation: -0.136, CI: -0.219 – -0.053, PE avg temp: -0.1184, CI: -0.216 – 0.021, PE seas temp: -0.101, CI: -0.271 – -0.069) (Fig. 3C). Again, the model containing all parameters was a better fit than the null (DIC full model: 789, DIC null model: 808) and also had a better fit than the interaction model or the model without the bat abundance parameter included (DIC interaction model: 791, DIC no bats: 799). For body size, the interaction between bat abundance and seasonal temperature variation is not significant (Fig 3D). Overall, the predictors explained less of the variance in body size (r^2^=0.72) than hindwing length. See code and outputs on Dryad.

## 4. Discussion

Combining species observations from iNaturalist and museum collections, a densely sampled *Actias* phylogeny (Fig. 1), and a comprehensive set of species distribution maps (SDMs) for 179 insectivorous bat species (Fig. 2), we investigated the relationship between hindwing tail length and biotic and abiotic drivers in the entirely tailed *Actias* + allies clade of Saturniidae (Fig 3). Our phylogeny captures 21 out of the approximately 40 *Actias* species and covers all major lineages, with only some species from known species complexes missing. To understand the evolutionary dynamics of this group, we estimated divergence times and used this dated phylogeny to infer species ancestral ranges. *Argema* + *Actias* diverged from their non-tailed sister taxa ∼20 mya and *Argema* and *Actias* diverged ∼15 mya, most likely when the lineage leading to *Argema* moved to Africa and the rest of *Actias* spread from the Indomalaya region (Fig. 2). Subsequently, *Actias* moved throughout the Eastern Palearctic (∼10 mya) and from Malaysia into the Indo-Australian archipelago (∼4mya). It is unclear when the European *Actias* (*Graellsia [Actias] isabellae*) and the North American *Actias* (*A. luna* and *truncatipennis*) colonized these regions respectively, although our analyses indicate that they both diverged from the rest of *Actias* ∼9 mya and sometime later become established in these areas (Fig. 1). Over the course of this time, the climate [81] and land masses were similar to current-day conditions, aside from movements of the Indo-Australian archipelago that continued until ∼5 mya (before *Actias* were present in this region) [46] and the closure of the Bering land bridge, which may have facilitated the movement of *Actias* to the North American continent [82,83]. The relatively young age of this clade is therefore one of the strengths of this study, as present-day patterns can be more reliably used to infer historic dynamics. Similarly, predation pressure has likely been relatively consistent throughout the evolution of *Actias.* Based on fossil evidence [84,85] and biogeographical reconstructions [84,86–89], large insectivorous bats had already become globally spread by this time (∼15 mya). Moreover, current dated bat phylogenies indicate that many extant lineages diversified 10-15 mya [85,90]. This rise in bat diversity and widespread prevalence of these predators could have made hindwing protrusions more profitable, as the night sky filled with more echolocators exploiting a greater depth of the prey community [91].

To estimate the effective pressure of bat predation, we built species distribution models (SDMs) for sufficiently large (10g or more) insectivorous bats and estimated abundance from these models (see Methods and Results sections for more details). We note that different bat species may exert differing predatory pressures on saturniid moths based on the specifics of their echolocation strategy or feeding guild [92], but given the large-scale nature of our data set and the generally similar diets of these aerial insectivores, we have considered insectivorous bats as a pooled group for the purposes of this study. To pit bat pressure against environmental factors (see Fig 3. for a list of predictors), we extracted values for these biotic and abiotic variables from hundreds of moth observations that we gathered from museum and community science specimens. Previous lab work has shown that tailed moths with ∼4cm difference in distance from body to tail tip can have a 25% increase in escape success from bats (Rubin et al. 2018). We therefore developed a method to extract a measure of absolute hindwing length from all moth photos, including those without a scale bar, which we believe will be helpful to future scientists interested in similar endeavors.

Analyzing these macroevolutionary data in a phylogenetically-informed linear mixed model framework provides evidence that bat predation pressure has likely exerted a selective force on the length of hindwing tails, while seasonal temperature variation has exerted a counterbalancing constraint on hindwing length (Fig. 3A). Moths with long tails are therefore less likely to be found in areas with fewer bats and more temperature fluctuation across the year. This result is supported by the positive association between the length of growing period and hindwing tail length. In essence, areas with longer periods of high plant productivity and more consistent temperature regimes appear to be more permissive of the evolution of long hindwing tails than areas with more restrictive seasons. Although weaker than the seasonality parameters, we found a negative association between hindwing length and average annual temperature and precipitation, indicating an opposite trend from Allen’s rule for endotherms, where appendages are expected to elongate in hotter, drier environments (as in [93]). This aligns with previous work indicating that Lepidoptera wings are not used for heat venting [94]. We did not find an effect of latitude in any of our models, indicating that the underlying drivers of wing trait evolution in this group are more complex than general latitudinal gradients. Additionally, while previous studies have found latitude to be an important correlate of bat diversity [95], others have found that it is not the most informative predictor, especially in the case of insectivores [96–98]. In congruence with this, we found relatively weak associations between insectivorous bat abundance and any of our climactic variables in the context of our global models (vif scores < 3; see code and outputs in Dryad). We did find an interaction effect between bat abundance and seasonal temperature variation in relation to tail length, however, indicating that bats and seasonality have their own relationship that influences tail length. That is, areas with less seasonal variability tend to host more bats as well as longer tailed moths (and vice versa, see Fig. S12 for an illustration of this interaction). From this interaction model, we also find that both seasonal temperature variation and bat abundance have their own appreciable effect on hindwing tail length. While the bat posterior distribution overlaps the zero line, there is a 0.91 probability that bat abundance is positively associated with moth tail length (Fig. 3B). Together, our analyses indicate that tails are locked in evolutionary tension between abiotic constraints and biotic pressure.

Contrary to our predictions, body size (forewing length) demonstrated an almost identical positive association with bat abundance as hindwing length (Fig. 3C, D). This could be because wing/body sizes are tightly integrated such that long hindwing tails require, or are made possible by, larger body sizes. This does not seem to be a ubiquitous rule in saturniids, as previous work in the subfamily Arsenurinae found an inverse relationship between hindwing length and body size (using forewing length as a proxy) [31]. However, arsenurine tails have a different structure than those of the Saturniinae (the subfamily containing *Actias*), in that they often protrude off the distal hindwing veins, rather than proximate ones [99]. We therefore think it is quite likely that these two subfamilies have different relationships between body size and hindwing tail length. In our clade of interest hindwing length scales with forewing length (R^2^ = 0.60) and thus most likely body size. Rather than simply being a necessary precursor for long hindwing tails, however, body size may be an anti-bat trait in itself. Bats seem to target prey relative to their own size, such that smaller bats eat smaller insects and larger bats are the main predators of large moths and beetles [100,101]. Whether this is due to handling, gape size, or echolocation limitations is still debated [53,102–104]. We found that the positive association between bat abundance and forewing length is complemented by a positive association between length of growing period and forewing length, again indicating that longer periods of forage availability allows for longer periods of larval feeding and larger adult body sizes [15]. These effects were to some extent countered by a negative association with precipitation. This may indicate a limitation on body size in regions with more rainfall, perhaps due to hampered foraging or increased larval mortality during bouts of heavy rain [105]. However, precipitation parameters from Worldclim should be considered with caution, especially from tropical regions with fewer climactic field data collection stations [106].

In addition to its use in our statistical models, comparative trait analyses revealed multiple origins of tail elongation but only one adaptive peak at the stem of the long-tailed moon moth clade, comprising all tailed species. This may be a result of the relatively limited number of species in this group and the strong phylogenetic signal underlying the tail trait. That is, while hindwing length varies considerably among these species, all species in this clade have tails, possibly making it more difficult to find the valleys between the morphological peaks [75,107]. Our results are congruent with a prior study that inferred a similar adaptive peak regime in *Argema + Actias* that was convergently repeated across the entire saturniid tree [12]. The multiple elongation and shrinkage events across our phylogeny indicates that the tail is a labile trait that could have become enhanced under conditions of high enough echolocating predator pressure and permissive environmental conditions, and that could relatively easily regress under more restrictive conditions. Previous research into the morphological lability of the fore- and hindwings of tailed swallowtail butterflies (Papillionidae), found similarly elevated hindwing shape diversity [108]. Lepidopteran wing shape variation is likely driven by different biological pressures on the two sets of wings, where forewings are essential for flight, while hindwings are helpful for maneuverability, but not entirely necessary [109–111]. Further, experimental evidence indicates that rather than being purely flight-driven, hindwings can play an important role in deflecting predators both during the day (in butterflies) [112] and night (in moths) [11,12].

While there are risks to making assumptions about past predator and prey dynamics based on extant forms, interactions, or distributions [113], the relative consistency of environmental conditions and bat presence strengthens our inferential power. Additionally, while our bat abundance estimates come with necessary assumptions and levels of uncertainty (e.g., species distribution models of extant species can be uncertain for species that are difficult to “observe”, as is the case with some insectivorous bats [114,115] and the population estimates were built from a global mammal dataset which could only provide coarse estimates [66]), we are ultimately interested in relative, rather than absolute, predator abundance. In general, species richness – the backbone upon which we built our abundance estimates – remains stable when ecological limits (most driven by climactic variables) are similar [116–118]. Thus, while extant bat distributions may not directly mirror historical ones, moths were clearly under intense selection pressure by echolocating bats in these regions.

In sum, results from this study, in conjunction with previous behavioral work [11,12] provide synergistic compelling evidence that predation pressure is associated with the elongation of hindwing tails in moon moths. Considering the absence of alternative selective forces (i.e., reproduction [1] or diurnal predation [14]) and the clear efficacy of short tails to increase escape success [12], we postulate that bat predation pressure drove the origins of the hindwing tail in Saturniidae. Hindwing tails with twisted and cupped ends have emerged five independent times across Saturniidae, three times in the Saturniinae (tribes: Saturniini, Attacini, Urotini/Bunaeini), once within the Arsenurinae [11,12,31], and once in Cercophaninae [19,119,120]. Phylogenetic inertia and the seemingly easily modifiable unit of wing imaginal discs in developing Lepidoptera [121] likely played a role in the evolution of tails. Contrary to the tail-elongating force of predation pressure, the elaboration of this trait appears to be limited by environmental factors. Indeed, the constraint of these abiotic variables may at times supersede the positive driver of predation. While developmental studies are needed to uncover the mechanism by which environment constrains tail enhancement (i.e., building a tail may require more nutritional resources and a longer growing season than building a more simplified hindwing), the negative association that we found between climatic variables, and the positive association with longer growing periods, provides evidence for an environmentally-mediated long-term cost of these appendages. A similar relationship was previously found between bright butterfly coloration, climatic variables, and bird diversity, indicating that trait elaboration of multiple kinds is likely limited by environmental factors [122]. Here, our study adds an important macroevolutionary lens to previous experimental predator-prey work. Uniting these two levels of information provides important advancement to our understanding of complex evolutionary dynamics and opens new lines of inquiry for future research [123]. Additional studies at an intermediate scale, testing the relationship between microhabitat, bat predation, and hindwing tails, could also reveal important detail about these dynamics. We emphasize the strength of multi-scale investigation for illuminating the relative pressures of competing eco-evolutionary forces that have shaped the origin and diversification of elaborate traits across taxonomic systems.

## Acknowledgments

Our sincere gratitude goes to Emily Ellis, who guided JJR in bioinformatics. David Plotkin helped with trouble-shooting phylogenetic and BioGeoBEARS analyses. Christian Couch and Amanda Markee contributed to DNA extraction and sequencing. Hailey Dansby collected photos of museum specimens. We appreciate David Grimaldi and Suzanne Green for help with AMNH specimen photos, as well as Stefan Naumann for providing *Actias felicis* photos and data. Sequenced specimens were collected with the help of several researchers, including Wayne Hsu, Charlie Mitter, and Richard Peigler. We thank José Miguel Ponciano for encouraging JJR to explore simulations at the start of this project, Emma Podietz for her help with ArcGIS, Sean Severud for help with figure design, and Michael Belitz for his support on species richness estimation.

## References

1. Rubin JJ, Kawahara AY. 2023 Sexual selection does not drive hindwing tail elaboration in a moon moth, Actias luna. Behavioral Ecology 34, 488–494. (doi:10.1093/beheco/arad019)

2. Graham ZA, Garde E, Heide-Jørgensen MP, Palaoro A V. 2020 The longer the better: Evidence that narwhal tusks are sexually selected. Biol Lett 16, 1–5. (doi:10.1098/rsbl.2019.0950)

3. Petrie M, Tim H, Carolyn S. 1991 Peahens prefer peacocks with elaborate trains. Anim Behav 41, 323–331. (doi:10.1016/S0003-3472(05)80484-1)

4. Crofts SB, Stankowich T. 2021 Stabbing spines: A review of the biomechanics and evolution of defensive spines. Integr Comp Biol 61, 655–667. (doi:10.1093/icb/icab099)

5. Garland T, Downs CJ, Ives AR. 2022 Trade-offs (And constraints) in organismal biology. Physiological and Biochemical Zoology 95, 82–112. (doi:10.1086/717897)

6. Basolo AL, Alcaraz G. 2003 The turn of the sword: Length increases male swimming costs in swordtails. Proceedings of the Royal Society B: Biological Sciences 270, 1631–1636. (doi:10.1098/rspb.2003.2388)

7. Somjee U. 2021 Positive allometry of sexually selected traits: Do metabolic maintenance costs play an important role? BioEssays 43, 1–13. (doi:10.1002/bies.202000183)

8. Thavarajah NK, Tickle PG, Nudds RL, Codd JR. 2016 The peacock train does not handicap cursorial locomotor performance. Sci Rep 6, 1–6. (doi:10.1038/srep36512)

9. Møller AP. 1996 The cost of secondary sexual characters and the evolution of cost-reducing traits. Ibis 138, 112–119. (doi:10.1111/j.1474-919x.1996.tb04317.x)

10. Scoble MJ. 1992 The Lepidoptera. Form, function and diversity. Oxford: Oxford University Press.

11. Barber JR, Leavell BC, Keener AL, Breinholt JW, Chadwell BA, McClure CJW, Hill GM, Kawahara AY. 2015 Moth tails divert bat attack: Evolution of acoustic deflection. Proceedings of the National Academy of Sciences 112, 2812–2816. (doi:10.1073/pnas.1421926112)

12. Rubin JJ, Hamilton CA, McClure CJW, Chadwell BA, Kawahara AY, Barber JR. 2018 The evolution of anti-bat sensory illusions in moths. Sci Adv 4, 1–10. (doi:10.1126/sciadv.aar7428)

13. Procheş Ş. 2005 The world’s biogeographical regions: Cluster analyses based on bat distributions. J Biogeogr 32, 607–614. (doi:10.1111/j.1365-2699.2004.01186.x)

14. Rubin JJ, Martin NW, Sieving KE, Kawahara AY. 2023 Testing bird-driven diurnal trade-offs of the moon moth’s anti-bat tail. Biol Lett 19, 1–5. (doi:10.1098/rsbl.2022.0428)

15. Nijhout HF, Riddiford LM, Mirth C, Shingleton AW, Suzuki Y, Callier V. 2014 The developmental control of size in insects. Wiley Interdiscip Rev Dev Biol 3, 113–134. (doi:10.1002/wdev.124)

16. McKenna KZ, Tao D, Nijhout HF. 2019 Exploring the role of insulin signaling in relative growth: A case study on wing-body scaling in Lepidoptera. Integr Comp Biol 59, 1324–1337. (doi:10.1093/icb/icz080)

17. Nijhout HF, Emlen DJ. 1998 Competition among body parts in the development and evolution of insect morphology. Proc Natl Acad Sci U S A 95, 3685–3689. (doi:10.1073/pnas.95.7.3685)

18. Macdonald WP, Martin A, Reed RD. 2010 Butterfly wings shaped by a molecular cookie cutter: Evolutionary radiation of lepidopteran wing shapes associated with a derived Cut/wingless wing margin boundary system. Evol Dev 12, 296–304. (doi:10.1111/j.1525-142X.2010.00415.x)

19. Hamilton CA, St Laurent RA, Dexter K, Kitching IJ, Breinholt JW, Zwick A, Timmermans MJTN, Barber JR, Kawahara AY. 2019 Phylogenomics resolves major relationships and reveals significant diversification rate shifts in the evolution of silk moths and relatives. BMC Evol Biol 19, 1–13. (doi:10.1186/s12862-019-1505-1)

20. Kawahara AY, Plotkin D, Espeland M, Meusemann K, Toussaint EFA. 2019 Phylogenomics reveals the evolutionary timing and pattern of butterflies and moths. PNAS, 1–7. (doi:10.1073/pnas.1907847116)

21. Tobler A, Nijhout HF. 2010 Developmental constraints on the evolution of wing-body allometry in Manduca sexta. Evol Dev 12, 592–600. (doi:10.1111/j.1525-142X.2010.00444.x)

22. Mousseau TA. 1997 Ectotherms follow the converse to Bergmann’s rule. 51, 630–632. (doi:10.2307/2411138)

23. Dmitriew CM. 2011 The evolution of growth trajectories: What limits growth rate? Biological Reviews 86, 97–116. (doi:10.1111/j.1469-185X.2010.00136.x)

24. Lehnert MS, Scriber JM, Gerard PD, Emmel TC. 2012 The ‘converse to Bergmann’s rule’ in tiger swallowtail butterflies: boundaries of species and subspecies wing traits are independent of thermal and host-plant induction. American Entomologist 53, 156–165.

25. Plaistow SJ, Tsuchida K, Tsubaki Y, Setsuda K. 2005 The effect of a seasonal time constraint on development time, body size, condition, and morph determination in the horned beetle *Allomyrina dichotoma* L. (Coleoptera: Scarabaeidae). Ecol Entomol 30, 692–699. (doi:10.1111/j.0307-6946.2005.00740.x)

26. Beerli N, Bärtschi F, Ballesteros-Mejia L, Kitching IJ, Beck J. 2019 How has the environment shaped geographical patterns of insect body sizes? A test of hypotheses using sphingid moths. J Biogeogr 46, 1687– 1698. (doi:10.1111/jbi.13583)

27. Brehm G, Zeuss D, Colwell RK. 2019 Moth body size increases with elevation along a complete tropical elevational gradient for two hyperdiverse clades. Ecography 42, 632–642. (doi:10.1111/ecog.03917)

28. Svensson EI, Gómez_Llano M, Waller JT. 2022 Out of the tropics: Macroevolutionary size trends in an old insect order are shaped by temperature and predators. J Biogeogr, 1–14. (doi:10.1111/jbi.14544)

29. Shelomi M, Zeuss D. 2017 Bergmann’s and Allen’s rules in native European and Mediterranean Phasmatodea. Front Ecol Evol 5, 1–13. (doi:10.3389/fevo.2017.00025)

30. D’Abrera B. 1995 Saturniidae Mundi. Antiquariat Goecke & Evers.

31. Hamilton CA, Winiger N, Rubin JJ, Breinholt J, Rougerie R, Kitching IJ, Barber JR, Kawahara AY. 2022 Hidden phylogenomic signal helps elucidate Arsenurine silkmoth phylogeny and the evolution of body size and wing shape trade-offs. Syst Biol 71, 859–874. (doi:10.1093/sysbio/syab090)

32. Miller WE. 1977 Wing Measure as a size index in Lepidoptera: the family Olethreutidae. Ann Entomol Soc Am 70, 253–256. (doi:10.1093/aesa/70.2.253)

33. Breinholt JW, Earl C, Lemmon AR, Moriarty Lemmon E, Xiao L, Kawahara AY. 2017 Resolving relationships among the megadiverse butterflies and moths with a novel pipeline for Anchored Phylogenomics. Syst Biol 0, 1–16. (doi:10.1093/sysbio/syx048)

34. Larsson A. 2014 AliView: A fast and lightweight alignment viewer and editor for large datasets. Bioinformatics 30, 3276–3278. (doi:10.1093/bioinformatics/btu531)

35. Kück P, Longo GC. 2014 FASconCAT-G: extensive functions for multiple sequence alignment preparations concerning phylogenetic studies. Front Zool 11, 81. (doi:10.1186/s12983-014-0081-x)

36. Nguyen LT, Schmidt HA, Von Haeseler A, Minh BQ. 2015 IQ-TREE: A fast and effective stochastic algorithm for estimating maximum-likelihood phylogenies. Mol Biol Evol 32, 268–274. (doi:10.1093/molbev/msu300)

37. Drummond AJ, Rambaut A. 2007 BEAST: Bayesian evolutionary analysis by sampling trees. BMC Evol Biol 7. (doi:10.1186/1471-2148-7-214)

38. Lanfear R, Calcott B, Ho SYW, Guindon S. 2012 PartitionFinder:Combined selection of partitioning schemes and substitution models for phylogenetic analyses. *UPB Scientific Bulletin*, Series B: Chemistry and Materials Science 29, 1695–1701. (doi:10.1093/molbev/mss020)

39. Mirarab S, Warnow T. 2015 ASTRAL-II: Coalescent-based species tree estimation with many hundreds of taxa and thousands of genes. Bioinformatics 31, i44–i52. (doi:10.1093/bioinformatics/btv234)

40. Drummond AJ, Suchard MA, Xie D, Rambaut A. 2012 Bayesian phylogenetics with BEAUti and the BEAST 1.7. Mol Biol Evol 29, 1969–1973. (doi:10.1093/molbev/mss075)

41. Maddison W, Maddison D. 2022 Mesquite: A modular system for evolutionary analysis. Version 3.60. http://www.mesquiteproject.org.

42. Gernhard T. 2008 The conditioned reconstructed process. J Theor Biol 253, 769–778. (doi:10.1016/j.jtbi.2008.04.005)

43. Matzke N. 2018 BioGeoBEARS: BioGeography with Bayesian (and likelihood) evolutionary analysis with R Scripts. *R Package*, Version 1.1.1. 1.

44. Matzke NJ. 2014 Model selection in historical biogeography reveals that founder-event speciation is a crucial process in island clades. Syst Biol 63, 951–970. (doi:10.1093/sysbio/syu056)

45. Toussaint EFA, Balke M. 2016 Historical biogeography of Polyura butterflies in the oriental Palaeotropics: trans-archipelagic routes and South Pacific island hopping. J Biogeogr 43, 1560–1572. (doi:10.1111/jbi.12741)

46. Lohman DJ, de Bruyn M, Page T, von Rintelen K, Hall R, Ng PKL, Shih H-T, Carvalho GR, von Rintelen T. 2011 Biogeography of the Indo-Australian Archipelago. Annu Rev Ecol Evol Syst 42, 205–226. (doi:10.1146/annurev-ecolsys-102710-145001)

47. Toussaint EFA et al. 2019 Out of the Orient: Post-Tethyan transoceanic and trans-Arabian routes fostered the spread of Baorini skippers in the Afrotropics. Syst Entomol 44, 926–938. (doi:10.1111/syen.12365)

48. Gladenkov AY, Oleinik AE, Marincovich L, Barinov KB. 2002 A refined age for the earliest opening of Bering Strait. Palaeogeogr Palaeoclimatol Palaeoecol 183, 321–328. (doi:10.1016/S0031-0182(02)00249-3)

49. Milne IR. 2006 Northern hemisphere plant disjunctions: A window on tertiary land bridges and climate change? Ann Bot 98, 465–472. (doi:10.1093/aob/mcl148)

50. Condamine FL, Rolland J, Morlon H. 2013 Macroevolutionary perspectives to environmental change. Ecol Lett 16, 72–85. (doi:10.1111/ele.12062)

51. Baker RJ, Bininda-Edmonds ORP, Mantilla-Meluk H, Porter CA, Van Den Bussche RA. 2012 Molecular time scale of diversification of feeding strategy and morphology in New World Leaf-Nosed Bats (Phyllostomidae): a phylogenetic perspective. In Evolutionary history of bats: fossils, molecules, and morphology (eds GF Gunnell, NB Simmons), pp. 385–409. New York: Cambridge University Press.

52. Simmons NB, Cirranello AL. In press. Bat species of the world: A taxonomic and geographic database, version 1.3. 2022.

53. Barclay RMR, Brigham RM. 1991 Prey detection, dietary niche breadth, and body size in bats: why are aerial insectivorous bats so small? Am Nat 137, 693–703. (doi:10.1080/0305006790150101)

54. Folk RA, Siniscalchi CM, Doby J, Kates HR, Manchester SR, Soltis PS, Soltis DE, Guralnick RP, Belitz M. 2023 Spatial phylogenetics of Fagales: Investigating the history of temperate forests. BioRxiv (doi:10.1101/2023.04.17.537249)

55. Abbott JC, Bota-Sierra CA, Guralnick R, Kalkman V, González-Soriano E, Novelo-Gutiérrez R, Bybee S, Ware J, Belitz MW. 2022 Diversity of Nearctic dragonflies and damselflies (Odonata). Diversity 14, 1–18. (doi:10.3390/d14070575)

56. Zizka A et al. 2019 CoordinateCleaner: Standardized cleaning of occurrence records from biological collection databases. Methods Ecol Evol 10, 744–751. (doi:10.1111/2041-210X.13152)

57. Herkt KMB, Barnikel G, Skidmore AK, Fahr J. 2016 A high-resolution model of bat diversity and endemism for continental Africa. Ecol Modell 320, 9–28. (doi:10.1016/j.ecolmodel.2015.09.009)

58. Smeraldo S et al. 2018 Ignoring seasonal changes in the ecological niche of non-migratory species may lead to biases in potential distribution models: lessons from bats. Biodivers Conserv 27, 2425–2441. (doi:10.1007/s10531-018-1545-7)

59. Hayes MA, Cryan PM, Wunder MB. 2015 Seasonally-dynamic presence-only species distribution models for a cryptic migratory bat impacted by wind energy development. PLoS One 10. (doi:10.1371/journal.pone.0132599)

60. McClure ML et al. 2021 A hybrid correlative-mechanistic approach for modeling winter distributions of North American bat species. J Biogeogr 48, 2429–2444. (doi:10.1111/jbi.14130)

61. Fick SE, Hijmans RJ. 2017 WorldClim 2: new 1-km spatial resolution climate surfaces for global land areas. International Journal of Climatology 37, 4302–4315. (doi:10.1002/joc.5086)

62. Amatulli G, Domisch S, Tuanmu M-N, Parmentier B, Ranipeta A, Malczyk J, Jetz W. 2018 A suite of global, cross-scale topographic variables for environmental and biodiversity modeling. Sci Data 5, 1–15. (doi:10.1038/sdata.2018.40)

63. Townsend J, DiMiceli C. 2015 MOD44B MODIS/Terra Vegetation Continuous Fields Yearly L3 Global 500m SIN Grid. NASA LP DAAC.

64. Kass JM et al. 2022 The global distribution of known and undiscovered ant biodiversity. Sci. Adv. 8.

65. Calabrese JM, Certain G, Kraan C, Dormann CF. 2014 Stacking species distribution models and adjusting bias by linking them to macroecological models. Global Ecology and Biogeography 23, 99–112. (doi:10.1111/geb.12102)

66. Greenspoon L et al. 2023 The global biomass of wild mammals. Proc Natl Acad Sci U S A 120. (doi:10.1073/pnas.2204892120)

67. Chamberlain S, Barve V, Mcglinn D, Oldoni D, Desmet P, Geffert L, Ram K. 2023 rgbif: Interface to the Global Biodiversity Information Facility API. R package version 3.3.7.

68. Rasband WS. 2018 ImageJ.

69. Acharya L. 1995 Sex-biased predation on moths by insectivorous bats. Anim Behav 49, 1461–1468. (doi:10.1016/0003-3472(95)90067-5)

70. Li D, Dinnage R, Nell LA, Helmus MR, Ives AR. 2020 phyr: An r package for phylogenetic species-distribution modelling in ecological communities. Methods Ecol Evol 11, 1455–1463. (doi:10.1111/2041-210X.13471)

71. Fischer G, van Velthuizen H, Nachtergaele FO. 2000 Global Agro-Ecological Zones Assessment: Methodology and Results.

72. Zuur AF, Ieno EN, Elphick CS. 2010 A protocol for data exploration to avoid common statistical problems. Methods Ecol Evol 1, 3–14. (doi:10.1111/j.2041-210x.2009.00001.x)

73. Revell LJ. 2012 phytools: An R package for phylogenetic comparative biology (and other things). Methods Ecol Evol 3, 217–223. (doi:10.1111/j.2041-210X.2011.00169.x)

74. Goolsby EW, Bruggeman J, Ané C. 2017 Rphylopars: fast multivariate phylogenetic comparative methods for missing data and within-species variation. Methods Ecol Evol 8, 22–27. (doi:10.1111/2041-210X.12612)

75. Ingram T, Mahler DL. 2013 SURFACE: Detecting convergent evolution from comparative data by fitting Ornstein-Uhlenbeck models with stepwise Akaike Information Criterion. Methods Ecol Evol 4, 416–425. (doi:10.1111/2041-210X.12034)

76. Kitching I, Rougerie R, Zwick A, Hamilton C, St Laurent R, Naumann S, Ballesteros Mejia L, Kawahara A. 2018 A global checklist of the Bombycoidea (Insecta: Lepidoptera). Biodivers Data J 6, 1–13. (doi:10.3897/BDJ.6.e22236)

77. Nässig WA. 1991 Biological observations and taxonomic notes on *Actias isabellae* (Graells) (Lepidoptera, Saturniidae). Nota lepidoptera 14, 131–143.

78. Ylla J, Peigler RS, Kawahara AY. 2005 Cladistic analysis of moon moths using morphology, molecules, and behaviour: Actias Leach, 1815; Argema Wallengren, 1858; Graellsia Grote, 1896 (Lepidoptera: Saturniidae). SHILAP Revista de Lepidoterologia 33, 299–317.

79. Ives AR. 2019 R2s for Correlated Data: Phylogenetic Models, LMMs, and GLMMs. Syst Biol 68, 234–251. (doi:10.1093/sysbio/syy060)

80. Miller WE. 1997 Diversity and evolution of tongue length in hawkmoths (Sphingidae). J Lepid Soc 51, 9– 31.

81. Steinthorsdottir M et al. 2021 The Miocene: The future of the past. Paleoceanogr Paleoclimatol 36, 1–71. (doi:10.1029/2020PA004037)

82. Hundsdoerfer AK, Kitching IJ, Wink M. 2005 A molecular phylogeny of the hawkmoth genus *Hyles* (Lepidoptera: Sphingidae, Macroglossinae). Mol Phylogenet Evol 35, 442–458. (doi:10.1016/j.ympev.2005.02.004)

83. Rubinoff D, Doorenweerd C. 2020 Systematics and biogeography reciprocally illuminate taxonomic revisions in the silkmoth genus *Saturnia* (Lepidoptera: Saturniidae). J Lepid Soc 74, 1–6. (doi:10.18473/lepi.74i1.a1)

84. Teeling EC, Springer MS, Madsen O, Bates P, O’brien SJ, Murphy WJ. 2005 A molecular phylogeny for bats illuminates biogeography and the fossil record. Science 307, 580–584. (doi:10.1126/science.1105113)

85. Shi JJ, Rabosky DL. 2015 Speciation dynamics during the global radiation of extant bats. Evolution 69, 1528–1545. (doi:10.1111/evo.12681)

86. Ruedi M, Friedli-Weyeneth N, Teeling EC, Puechmaille SJ, Goodman SM. 2012 Biogeography of Old World emballonurine bats (Chiroptera: Emballonuridae) inferred with mitochondrial and nuclear DNA. Mol Phylogenet Evol 64, 204–211. (doi:10.1016/j.ympev.2012.03.019)

87. Lamb JM et al. 2008 Phylogeography and predicted distribution of African-Arabian and Malagasy populations of giant mastiff bats, *Otomops* spp. (Chiroptera: Molossidae). Acta Chiropt 10, 21–40. (doi:10.3161/150811008X331063)

88. HoráČek I, Fejfar O, Hulva P. 2006 A new genus of vespertilionid bat from Early Miocene of Jebel Zelten, Libya, with comments on Scotophilus and early history of vespertilionid bats (Chiroptera). Lynx 37, 131– 150.

89. Chornelia A, Hughes AC. 2022 The evolutionary history and ancestral biogeographic range estimation of old-world Rhinolophidae and Hipposideridae (Chiroptera). BMC Ecol Evol 22. (doi:10.1186/s12862-022-02066-x)

90. Upham NS, Esselstyn JA, Jetz W. 2019 Inferring the mammal tree: Species-level sets of phylogenies for questions in ecology, evolution, and conservation. PLoS Biol 17, e3000494. (doi:10.1371/journal.pbio.3000494)

91. Alberdi A et al. 2020 DNA metabarcoding and spatial modelling link diet diversification with distribution homogeneity in European bats. Nat Commun 11, 1–8. (doi:10.1038/s41467-020-14961-2)

92. Denzinger A, Schnitzler HU. 2013 Bat guilds, a concept to classify the highly diverse foraging and echolocation behaviors of microchiropteran bats. Front Physiol 4, 1–15. (doi:10.3389/fphys.2013.00164)

93. Alhajeri BH, Fourcade Y, Upham NS, Alhaddad H. 2020 A global test of Allen’s rule in rodents. Global Ecology and Biogeography 29, 2248–2260. (doi:10.1111/geb.13198)

94. Wasserthal LT. 1975 The role of butterfly wings in regulation of body temperature. J Insect Physiol 21, 1921–1930. (doi:10.1016/0022-1910(75)90224-3)

95. Kaufman DM, Willig MR. 1998 Latitudinal patterns of mammalian species richness in the New World: The effects of sampling method and faunal group. J Biogeogr 25, 795–805. (doi:10.1046/j.1365-2699.1998.2540795.x)

96. Martins MA, Carvalho WD De, Dias D, Franca DDS, Oliveira MBD, Peracchi AL. 2015 Bat species richness (Mammalia, Chiroptera) along an elevational gradient in the Atlantic forest of southeastern Brazil. Acta Chiropt 17, 401–409. (doi:10.3161/15081109ACC2015.17.2.016)

97. Bogoni JA, Carvalho-Rocha V, Ferraz KMPMB, Peres CA. 2021 Interacting elevational and latitudinal gradients determine bat diversity and distribution across the Neotropics. Journal of Animal Ecology. (doi:10.1111/1365-2656.13594)

98. Patten MA. 2004 Correlates of species richness in North American bat families. J Biogeogr 31, 975–985. (doi:10.1111/j.1365-2699.2004.01087.x)

99. Zhong M, Hilla GM, Gomez JP, Plotkin D, Barber JR, Kawahara AY. 2016 Quantifying wing shape and size of saturniid moths with geometric morphometrics. J Lepid Soc 70, 99–107. (doi:10.18473/lepi.70i2.a4)

100. Wray AK, Peery MZ, Jusino MA, Kochanski JM, Banik MT, Palmer JM, Lindner DL, Gratton C. 2021 Predator preferences shape the diets of arthropodivorous bats more than quantitative local prey abundance. Mol Ecol 30, 855–873. (doi:10.1111/mec.15769)

101. Jones G. 1990 Prey selection by the greater horseshoe bat (*Rhinolophus ferrumequinum*): Optimal foraging by echolocation? Journal of Animal Ecology 59, 587–602.

102. Gonsalves L, Bicknell B, Law B, Webb C, Monamy V. 2013 Mosquito consumption by insectivorous bats: Does size matter? PLoS One 8. (doi:10.1371/journal.pone.0077183)

103. Giacomini G, Herrel A, Chaverri G, Brown RP, Russo D, Scaravelli D, Meloro C. 2022 Functional correlates of skull shape in Chiroptera: feeding and echolocation adaptations. Integr Zool 17, 430–442. (doi:10.1111/1749-4877.12564)

104. Bogdanowicz W, Fenton MB, Daleszczyk K. 1999 The relationships between echolocation calls, morphology and diet in insectivorous bats. J Zool 247, 381–393. (doi:10.1111/j.1469-7998.1999.tb01001.x)

105. Kamata N, Igarashi Y. 1994 Influence of rainfall on feeding behavior, growth, and mortality of larvae of the beech caterpillar, Quadricalcarifera punctatella (Motschulsky) (Lep., Notodontidae). Journal of Applied Entomology 118, 347–353. (doi:10.1111/j.1439-0418.1994.tb00810.x)

106. Soria-Auza RW, Kessler M, Bach K, Barajas-Barbosa PM, Lehnert M, Herzog SK, Böhner J. 2010 Impact of the quality of climate models for modelling species occurrences in countries with poor climatic documentation: a case study from Bolivia. Ecol Modell 221, 1221–1229. (doi:10.1016/j.ecolmodel.2010.01.004)

107. Graham CH, Storch D, Machac A. 2018 Phylogenetic scale in ecology and evolution. Global Ecology and Biogeography 27, 175–187. (doi:10.1111/geb.12686)

108. Owens HL, Lewis DS, Condamine FL, Kawahara AY, Guralnick RP. 2020 Comparative phylogenetics of Papilio butterfly wing shape and size demonstrates independent hindwing and forewing evolution. Syst Biol 69, 813–819. (doi:10.1093/sysbio/syaa029)

109. Jantzen B, Eisner T. 2008 Hindwings are unnecessary for flight but essential for execution of normal evasive flight in Lepidoptera. Proc Natl Acad Sci U S A 105, 16636–40. (doi:10.1073/pnas.0807223105)

110. Stylman M, Penz CM, DeVries P. 2020 Large hind wings enhance gliding performance in ground effect in a Neotropical butterfly (Lepidoptera: Nymphalidae). Ann Entomol Soc Am 113, 15–22. (doi:10.1093/aesa/saz042)

111. Le Roy C, Cornette R, Llaurens V, Debat V. 2019 Effects of natural wing damage on flight performance in Morpho butterflies: what can it tell us about wing shape evolution? J Exp Biol 222, jeb204057. (doi:10.1242/jeb.204057)

112. Chotard A, Ledamoisel J, Decamps T, Herrel A, Chaine AS, Llaurens V, Debat V. 2022 Evidence of attack deflection suggests adaptive evolution of wing tails in butterflies. Proceedings of the Royal Society B: Biological Sciences 289. (doi:10.1098/rspb.2022.0562)

113. Losos JB. 2011 Convergence, adaptation, and constraint. Evolution 65, 1827–1840. (doi:10.1111/j.1558-5646.2011.01289.x)

114. Razgour O, Rebelo H, Di Febbraro M, Russo D. 2016 Painting maps with bats: species distribution modelling in bat research and conservation. Hystrix 27, 1–8. (doi:10.4404/hystrix-27.1-11753)

115. Delgado-Jaramillo M, Aguiar LMS, Machado RB, Bernard E. 2020 Assessing the distribution of a species-rich group in a continental-sized megadiverse country: Bats in Brazil. Divers Distrib 26, 632–643. (doi:10.1111/ddi.13043)

116. Rabosky DL, Hurlbert AH. 2015 Species richness at continental scales is dominated by ecological limits. American Naturalist 185, 572–583. (doi:10.1086/680850)

117. Hawkins BA et al. 2012 Different evolutionary histories underlie congruent species richness gradients of birds and mammals. J Biogeogr 39, 825–841. (doi:10.1111/j.1365-2699.2011.02655.x)

118. Dornelas M, Gotelli NJ, McGill B, Shimadzu H, Moyes F, Sievers C, Magurran AE. 2014 Assemblage time series reveal biodiversity change but not systematic loss. Science (1979) **344**, 296–299. (doi:10.1126/science.1248484)

119. Wolfe KL, Balcázar-Lara MA. 1994 Chile’s *Cercophana venusta* and its immature stages (Lepidoptera: Cercophanidae). Tropical Lepdioptera 5, 35–42.

120. Aiello BR et al. 2021 Adaptive shifts underlie the divergence in wing morphology in bombycoid moths. Proceedings of the Royal Society B: Biological Sciences 288, 20210677. (doi:10.1098/rspb.2021.0677)

121. Fisher CR, Wegrzyn JL, Jockusch EL. 2020 Co-option of wing-patterning genes underlies the evolution of the treehopper helmet. Nat Ecol Evol 4, 250–260. (doi:10.1038/s41559-019-1054-4)

122. Dalrymple RL, Flores-Moreno H, Kemp DJ, White TE, Laffan SW, Hemmings FA, Hitchcock TD, Moles AT. 2018 Abiotic and biotic predictors of macroecological patterns in bird and butterfly coloration. Ecol Monogr 88, 204–224. (doi:10.1002/ecm.1287)

123. McCleery R et al. 2023 Uniting experiments and big data to advance ecology and conservation. Trends Ecol Evol, 1–10. (doi:10.1016/j.tree.2023.05.010)

